# Chromosome engineering to correct a complex rearrangement on Chromosome 8 reveals the effects of 8p syndrome on gene expression and neural differentiation

**DOI:** 10.1101/2024.11.17.624023

**Authors:** Sophia N. Lee, Erin C. Banda, Lu Qiao, Sarah L. Thompson, Karan Singh, Ryan A. Hagenson, Teresa Davoli, Stefan F. Pinter, Jason M. Sheltzer

## Abstract

Chromosomal rearrangements on the short arm of Chromosome 8 cause 8p syndrome, a rare developmental disorder characterized by neurodevelopmental delays, epilepsy, and cardiac abnormalities. While significant progress has been made in managing the symptoms of 8p syndrome and other conditions caused by large-scale chromosomal aneuploidies, no therapeutic approach has yet been demonstrated to target the underlying disease-causing chromosome. Here, we establish a two-step approach to eliminate the abnormal copy of Chromosome 8 and restore euploidy in cells derived from an individual with a complex rearrangement of Chromosome 8p. Transcriptomic analysis revealed 361 differentially expressed genes between the proband and the euploid revertant, highlighting genes both within and outside the 8p region that may contribute to 8p syndrome pathology. Furthermore, we demonstrate that the proband exhibits a significant defect in neural differentiation that could be partially rescued by treatment with small-molecule inhibitors of cell death. Our work demonstrates the feasibility of using chromosome engineering to correct complex aneuploidies in vitro and establishes a platform to further dissect the pathophysiology of 8p syndrome and other conditions caused by chromosomal rearrangements.

## Introduction

Large-scale gene copy number alterations are the most common cause of miscarriage and developmental disability in humans (Sheltzer and Amon 2011). Disorders like Down syndrome, caused by the triplication of Chromosome 21, and Cri du chat syndrome, caused by the deletion of a region of Chromosome 5p, affect the expression of hundreds of genes and perturb normal development. 8p syndrome, caused by rearrangements affecting the p arm of Chromosome 8, represents a rare class of developmental disorders and impacts about 1 in 10,000-30,000 live births a year (Lo Bianco et al. 2020).

Several different types of 8p rearrangements have been documented in the medical literature. The most common alteration affecting this region consists of a terminal distal deletion and an inverted duplication proximal to the Chromosome 8 centromere, termed invdupdel(8p) (Santucci et al. 2024). This rearrangement arises from aberrant pairing and recombination events during meiosis, which may be due to the presence of the ß-defensin repeats and olfactory gene receptor clusters found within this region (Giglio et al. 2001). Individuals affected by 8p syndrome present with a variety of clinical phenotypes, including agenesis of the corpus callosum, epilepsy, and other neurodevelopmental delays (Vibert et al. 2022).

Over the past several decades, the medical management of individuals with copy number disorders like Down syndrome has improved remarkably (Hendrix et al. 2021; Skotko et al. 2013),. Aggressive and early interventions designed to address drivers of morbidity in these populations have resulted in the average lifespan of an individual with Down syndrome increasing from 25 years in 1983 to around 60 years in 2010 (Glasson et al. 2016). However, to date, the successful treatment of individuals with copy number disorders has relied almost entirely on symptom management. Therapies designed to address the underlying genetic disorder, for instance, by inhibiting the function of a dosage-sensitive protein encoded in a triplicated region, have not proven to be successful (Jiang et al. 2013). The failure of targeted therapies to produce clinical benefit raises the possibility that other interventions may best be able to correct the underlying genetic pathology.

An alternate approach to treat disorders caused by large-scale chromosomal abnormalities would be to directly manipulate the causative chromosome(s). For these techniques, broadly called “chromosome therapies”, the disease-causing chromosome may be silenced, eliminated, or replaced with a non-pathological copy (Gupta et al. 2024; Paulis et al. 2015; Zuo et al. 2017). Pioneering work conducted in iPS cells derived from individuals with Down syndrome demonstrated that the extra copy of Chromosome 21 could be silenced through the use of the XIST noncoding RNA or removed through the introduction of a genetic negative selection cassette. Silencing or eliminating the extra chromosome reverted many pathological alterations induced by the trisomy, including defects in neural differentiation, (Bansal et al.; Gupta et al. 2024; Li et al. 2012). Further research revealed that the generation and passaging of iPS cells themselves could promote reversion to a euploid state, likely because of the detrimental fitness effects of the initial aneuploidy (Akutsu et al. 2022; Ya et al. 2024). “Chromosome therapy” may be particularly appealing for addressing a complex disorder like 8p syndrome, as no individual dosage-sensitive gene within this region has been demonstrated to drive pathological development, and individuals with this disorder exhibit significant variability in the size and location of the chromosomal rearrangements (Lo Bianco et al. 2020; Okur et al. 2021; Vibert et al. 2022),

A number of technical hurdles remain for implementing chromosome therapy as a medical intervention (Gupta et al. 2024). However, while this approach has not yet been successfully translated into the clinic, chromosome therapy can be utilized to generate a variety of powerful research tools. In particular, chromosome therapy can be applied to create matched cells that have or lack a specific chromosome abnormality but are otherwise genetically identical. Such cell line pairs bypass the confounding effects of genetic variability that complicate comparisons between cells derived from an individual with a chromosomal abnormality and their unaffected siblings or parents. Isogenic cell pairs can be ideal tools for drug screens, gene expression analysis, and investigations into the biological consequences of a specific chromosome rearrangement.

To date, successful examples of chromosome therapy applied *in vitro* have been limited to trisomies and ring chromosomes, (Akutsu et al. 2022; Bershteyn et al. 2014; Hirota et al. 2017; Inoue et al. 2019; Li et al. 2012, 2017). To our knowledge, the elimination of a complex rearrangement like invdupdel(8p) has not been reported. Here, we leveraged cutting-edge chromosome engineering techniques to derive a euploid revertant cell line from iPS cells established from an individual with invdupdel(8p). We then applied this matched pair of isogenic cell lines to interrogate the effects of invdupdel(8p) on gene expression and neural differentiation.

## Results

### Whole-genome sequencing of invdupdel(8p) iPS cells

8p syndrome is characterized by a wide spectrum of clinical phenotypes, including neurological, visual, and musculoskeletal abnormalities (Figure 1A). We received a line of iPS cells with invdupdel(8p) from Project 8p, a philanthropic group dedicated to improving the care of individuals affected by 8p syndrome. We performed low pass whole-genome sequencing to determine the extent of the copy number alterations exhibited by the proband. The deletion spanned from the telomere of Chromosome 8p to position 7,247,573 (∼7.2 Mb) and encompassed 23 protein-coding genes, while the duplication spanned from position 11,828,865 – 40,361,770 (∼28.5 Mb) and encompassed 172 protein-coding genes (Figure 1B-C) (Durinck et al. 2005, 2009; Harrison et al. 2024). No other aneuploidies were detected in the proband. A prior study of 8p syndrome reported a median deletion size of 7 Mb and a median duplication size of 25.8 Mb, indicating that the magnitude of the alterations in this proband are typical for this condition (Okur et al. 2021).

**Figure 1:**
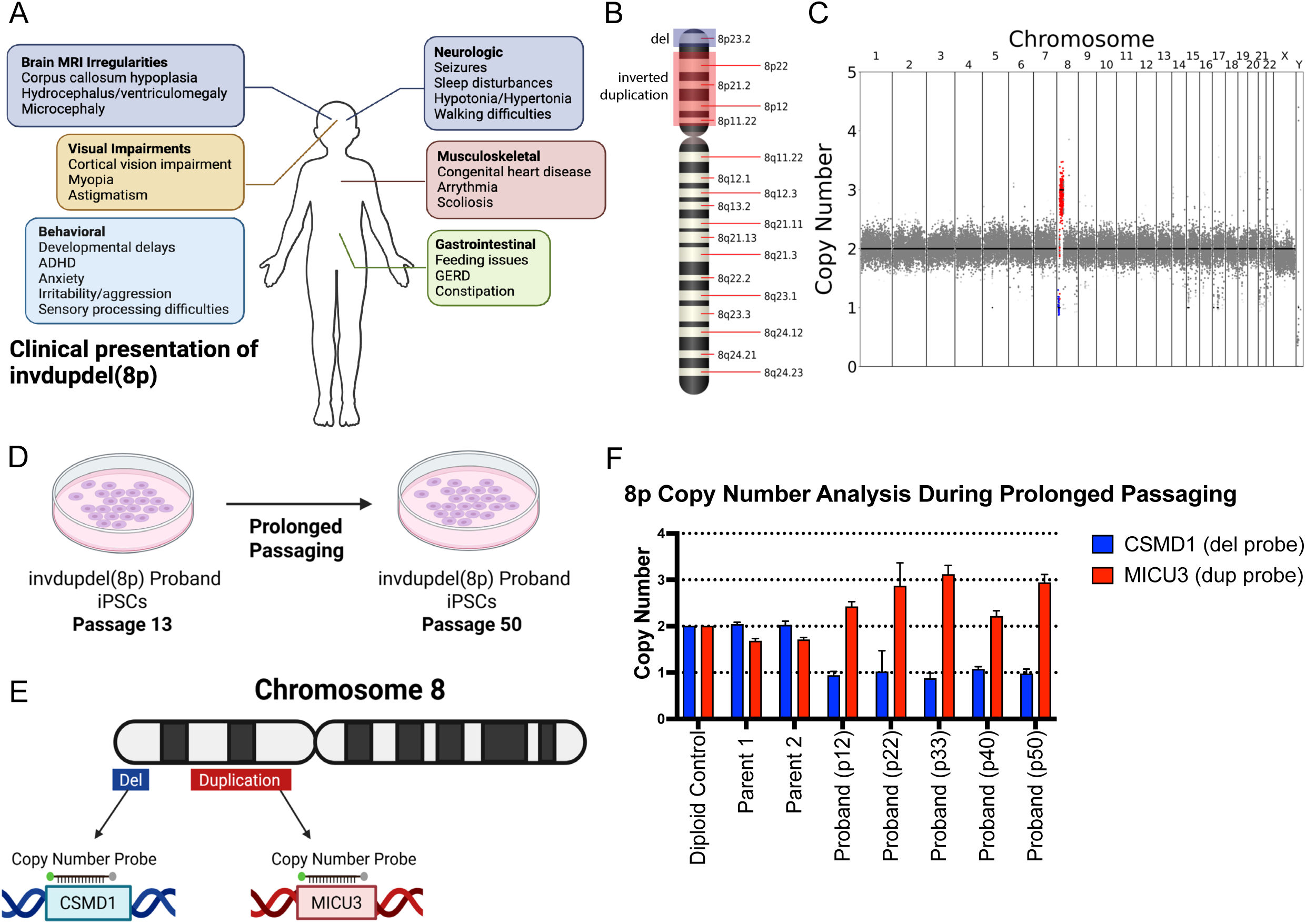
Karyotype of the invdupdel(8p) proband iPS line A. Schematic listing commonly observed symptoms in individuals affected by invdupdel(8p). B. Diagram delineating altered regions in the proband. The deleted region is marked in blue. The region that is inverted and duplicated is marked in red. C. Karyotype of the invdupdel(8p) proband iPS line. The deleted region is highlighted in blue. The region that is inverted and duplicated is marked in red. D. Schematic detailing the prolonged passaging of proband cells to induce spontaneous loss of the rearranged copy of Chromosome 8. E. Generation of copy number TaqMan probes specific to genes contained within the invdupdel(8p) altered regions. *CSMD1* is a gene located within the deleted region, and *MICU3* is a gene located within the duplicated region. F. Copy numbers of *CSMD1* and *MICU3* after passaging proband cells for 12, 22, 33, 40, and 50 passages. Mean ± SEM, data from representative trials are shown (n ≥ 3 total trials). Genomic DNA from the near-diploid MCF10A cell line was used as a control.

### Prolonged passaging of invdupdel(8p) iPS cells fails to restore euploidy

Previous reports in the literature have demonstrated that the prolonged growth of aneuploid iPS cells in culture can sometimes result in the isolation of derivative cells with euploid karyotypes, (Bershteyn et al. 2014; Ya et al. 2024). This may be because most aneuploidies confer a fitness disadvantage in primary cells, and rare euploid cells arising from chromosome missegregation events are able to outcompete the aneuploid starting population and overtake the culture (Thompson and Compton 2010; Vasudevan et al. 2021). We therefore sought to determine whether we could isolate invdupdel(8p) revertants with a euploid karyotype through continuous passaging (Figure 1D). We grew the proband’s cells for approximately 130 days in culture, until passage 50. We then designed a TaqMan-based strategy to assess the copy number of the invdupdel(8p) region (Figure 1E) (Mayo et al. 2010). qPCR confirmed the presence of two copies of the proband-deleted region and two copies of the proband-duplicated region in iPS cells derived from the proband’s parents (Figure 1F). We performed qPCR on the proband cell line at five timepoints spanning from passage 12 to passage 50, but we did not detect any evidence for the emergence of a euploid population (Figure 1F). We conclude that prolonged passaging is insufficient to restore euploidy in this proband’s iPS cells.

### Inducing missegregation to generate iPS cells with trisomy 8

We noted that previous reports describing the successful isolation of euploid revertants from aneuploid iPS cell lines were primarily obtained from trisomic starting populations, in which a single chromosome missegregation event generated cells with the desired disomic karyotype. In contrast, isolation of euploid revertants from invdupdel(8p) cells would require both the loss of the chromosome harboring the rearrangement and duplication of the remaining homologue. We therefore hypothesized that two possible factors could explain our failure to isolate spontaneous revertants from invdupdel(8p) iPS cells. First, the rate of chromosome missegregation in this iPS cell line may not have been sufficient to produce the two missegregation events necessary to restore euploidy. Alternately, monosomies are very poorly tolerated by primary cells, and even transient loss of the rearranged copy of 8p may have been incompatible with cellular viability, (Biancotti et al. 2012; Chunduri et al. 2021; Magnuson et al. 1985).

To circumvent these challenges, we developed an alternate approach to facilitate the isolation of invdupdel(8p) revertant cells. We treated the proband cells with a sublethal dose of the TTK (MPS1) inhibitor AZ3146, which we and others have shown ablates the mitotic checkpoint and enhances the rate of chromosome missegregation,(Hewitt et al. 2010; Kwiatkowski et al. 2010; Santaguida et al. 2010; Vasudevan et al. 2020). We hypothesized that transient exposure to AZ3146 could have two potential benefits: first, by increasing the frequency of chromosome missegregation, we could have a greater likelihood of isolating cells that spontaneously acquired a euploid karyotype. Alternately, treatment with AZ3146 could lead to the incidental production of cells trisomic for Chromosome 8, which could then serve as a stable intermediate for the subsequent isolation of a euploid revertant cell line (Figure 2A).

**Figure 2:**
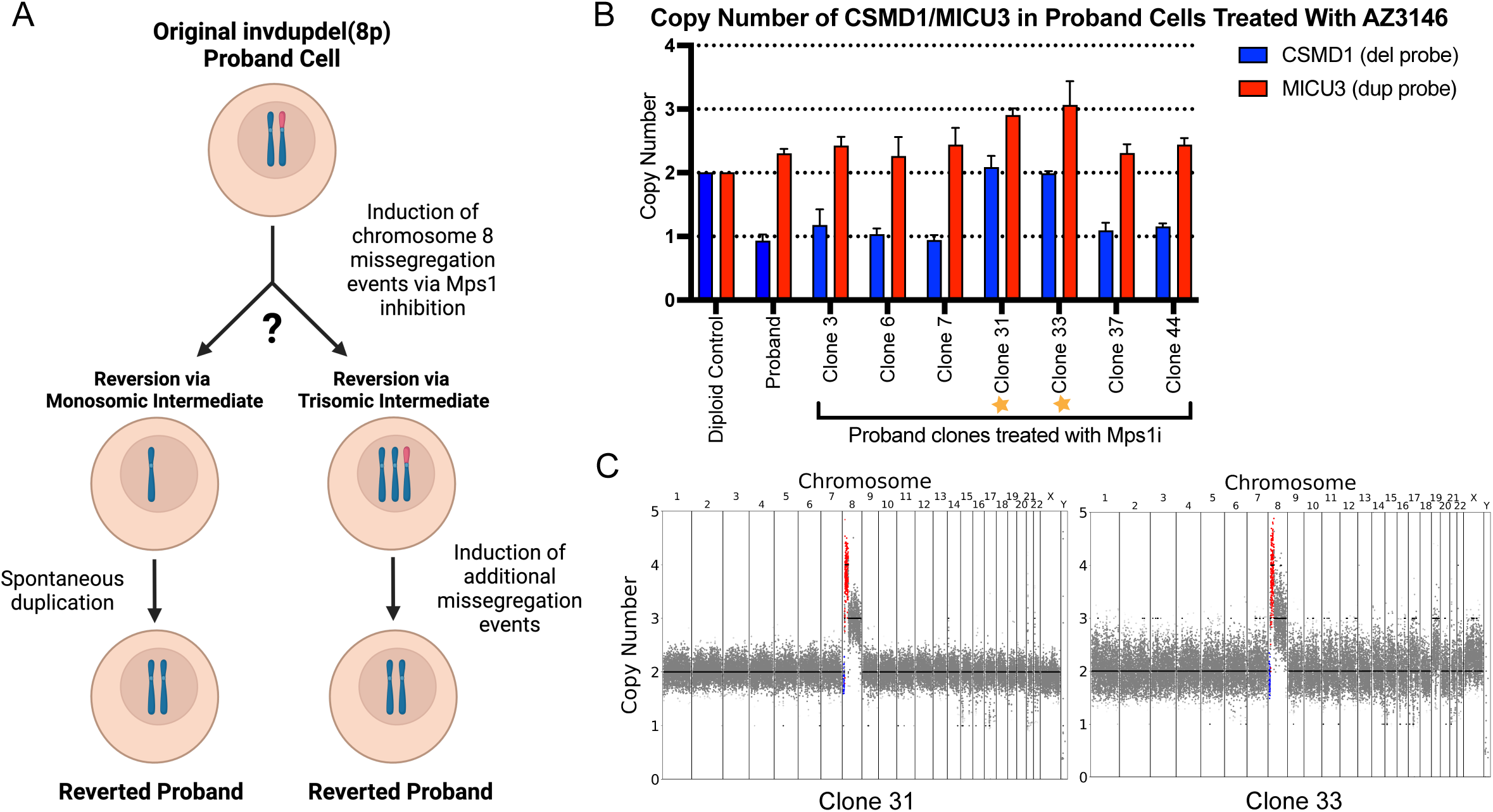
Generation of a Chromosome 8 trisomy through TTK inhibition A. Schematic describing two possible ways by which euploidy could be restored in proband cells after treatment with AZ3146. One method consists of the loss of the rearranged copy of Chromosome 8 coupled with the duplication of the remaining chromosome. The second method consists of gaining an unrearranged copy of Chromosome 8 and then subsequently losing the rearranged homologue. B. Copy numbers of *CSMD1* and *MICU3* in proband clones treated with AZ3146. Trisomic clones 31 and 33 are labeled with stars. Mean ± SEM, data from representative trials are shown (n ≥ 3 total trials). Genomic DNA from MCF10A cells was used as a diploid control. C. Karyotypes of two clones that have gained an additional unrearranged copy of Chromosome 8 following AZ3146 treatment.

Following treatment with AZ3146, we isolated 52 viable iPS clones. TaqMan qPCR using 8p-specific probes did not reveal any clones with a euploid karyotype. However, we identified two clones that appeared to have acquired a trisomy of Chromosome 8 (Figure 2B and S1). We performed low pass whole-genome sequencing on these clones and we established that they harbored a trisomy of Chromosome 8q, a tetrasomy of the duplicated region of Chromosome 8p, and a disomy of the 8p-deleted region. Clone 33 also exhibited a partial gain of Chromosome 19, while no other aneuploidies were apparent in clone 31 (Figure 2C). We therefore chose to use clone 31 in our subsequent efforts to restore Chromosome 8 euploidy.

### Inducing targeted missegregation to generate a euploid revertant iPS cell line

Simultaneous to our efforts to accelerate chromosome missegregation with an TTK inhibitor, Davoli and colleagues published a new system for targeted chromosome missegregation (Bosco et al. 2023). In this approach, called KaryoCreate, catalytically-inactive dCas9 is used to target a checkpoint-deficient kinetochore protein to a specific mammalian centromere (Figure 3A). The mutant kinetochore protein prevents correct amphitelic biorientation, leading to the missegregation of the targeted chromosome. We therefore sought to apply the KaryoCreate system to restore euploidy in our invdupdel(8p) iPS cell lines. We elected to conduct these experiments using clone 31, which harbors a trisomy of Chromosome 8, as inducing loss of the rearranged copy of Chromosome 8 would be sufficient to restore euploidy in this clone.

**Figure 3:**
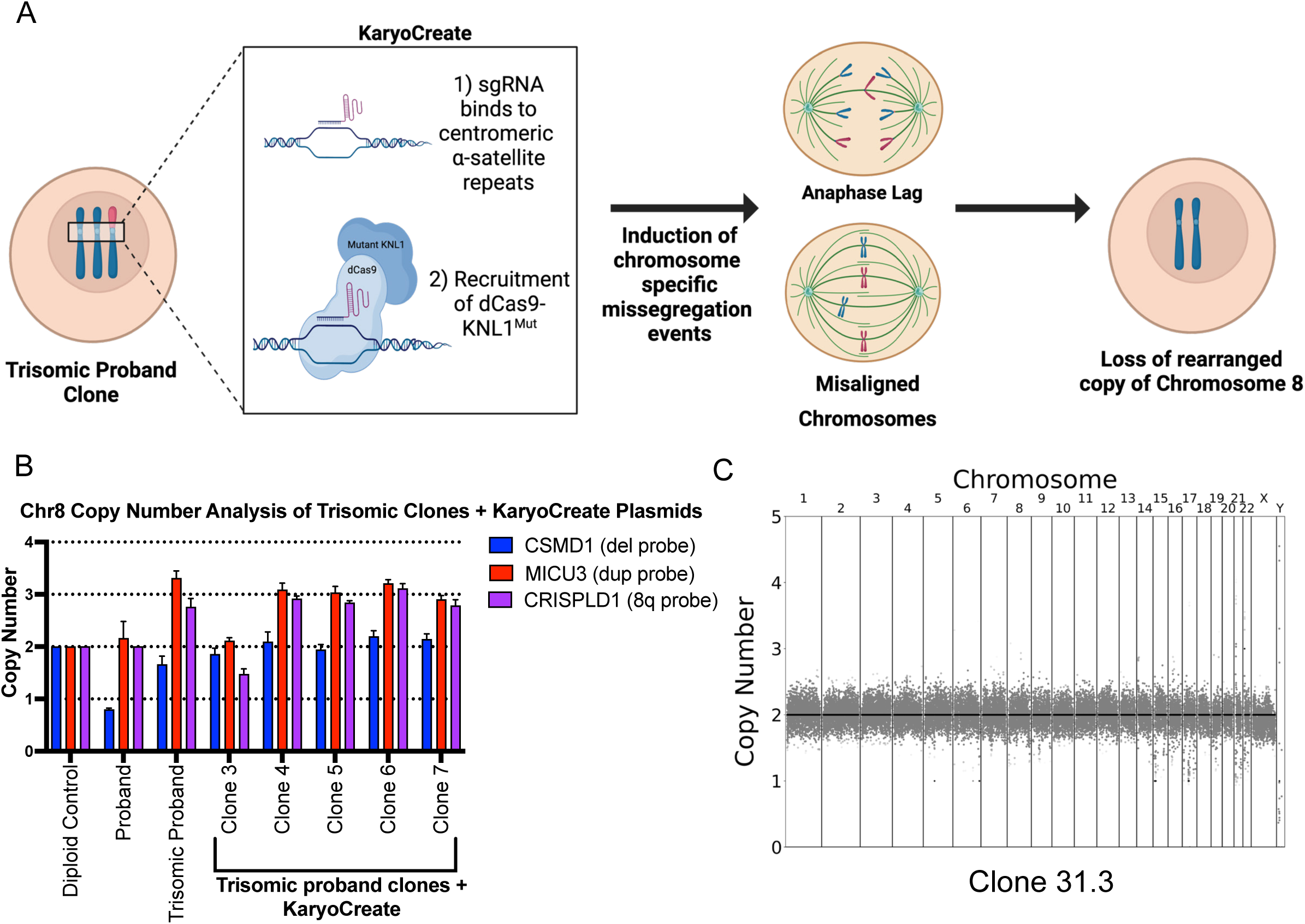
Restoration of euploidy using the KaryoCreate system A. Schematic describing the use of the KaryoCreate system to induce the missegregation of a specific chromosome. B. Copy numbers of *CSMD1*, *MICU3*, and *CRISPLD1* (a gene located on the q arm of Chromosome 8), in trisomic proband clones that were transiently transfected with KaryoCreate plasmids. Mean ± SEM, data from representative trials are shown (n ≥ 3 total trials). Genomic DNA extracted from MCF10A cells was used as a diploid control. C. Karyotype of the euploid revertant clone 31.3.

We transiently transfected clone 31 with plasmids encoding the dCas9 fusion protein and a gRNA specific to the Chromosome 8 centromere. We then isolated 43 viable clones and screened them via TaqMan to assess the copy number of Chromosome 8. qPCR analysis identified one clone that appeared to have lost the Chromosome 8 trisomy (Figure 3B and S2A). To rule out the possibility that this clone resulted from cell culture contamination, we performed STR profiling on iPS cells from the two parents, the proband, and the apparent revertant. This analysis confirmed that the revertant clone was derived from the proband, and analysis of the D8S1179 marker on Chromosome 8 revealed that it harbored an isodisomy for the paternal copy of Chromosome 8 (Figure S2B). Next, we performed low pass whole-genome sequencing on the revertant which confirmed the loss of the rearranged chromosome and the restoration of a euploid karyotype (Figure 3C). We compared proliferation between the proband and the revertant clones and we found that the revertant cells divided moderately more rapidly than the proband line, consistent with the 8p aneuploidy exerting a negative effect on cellular fitness (Figure S2C). The minor overall difference in proliferation rates may partially explain why prolonged passaging was insufficient to restore euploidy in the proband (Figure 1F).

### Gene expression analysis in proband and revertant iPS cells

To improve our understanding of how invdupdel(8p) affects normal development, we performed RNA-seq on iPS cells derived from the proband, the two parents, and on the revertant clone. We first compared gene expression on Chromosome 8 between the proband and the revertant (Figure 4A). In the region corresponding to the deletion, genes in the proband were expressed on average 43% lower, and in the region corresponding to the duplication, genes in the proband were expressed on average 56% higher, relative to the revertant. We also noted a region on the q arm of Chromosome 8 that exhibited a moderate increase in gene expression in the proband, but our whole-genome sequencing (Figure 1C and 3C) as well as TaqMan using a probe that recognizes this region (Figure S3A) confirmed that 8q remained diploid. We did not observe any chromosome-specific changes in gene expression along other chromosomes in the proband (Figure S3B-C).

**Figure 4:**
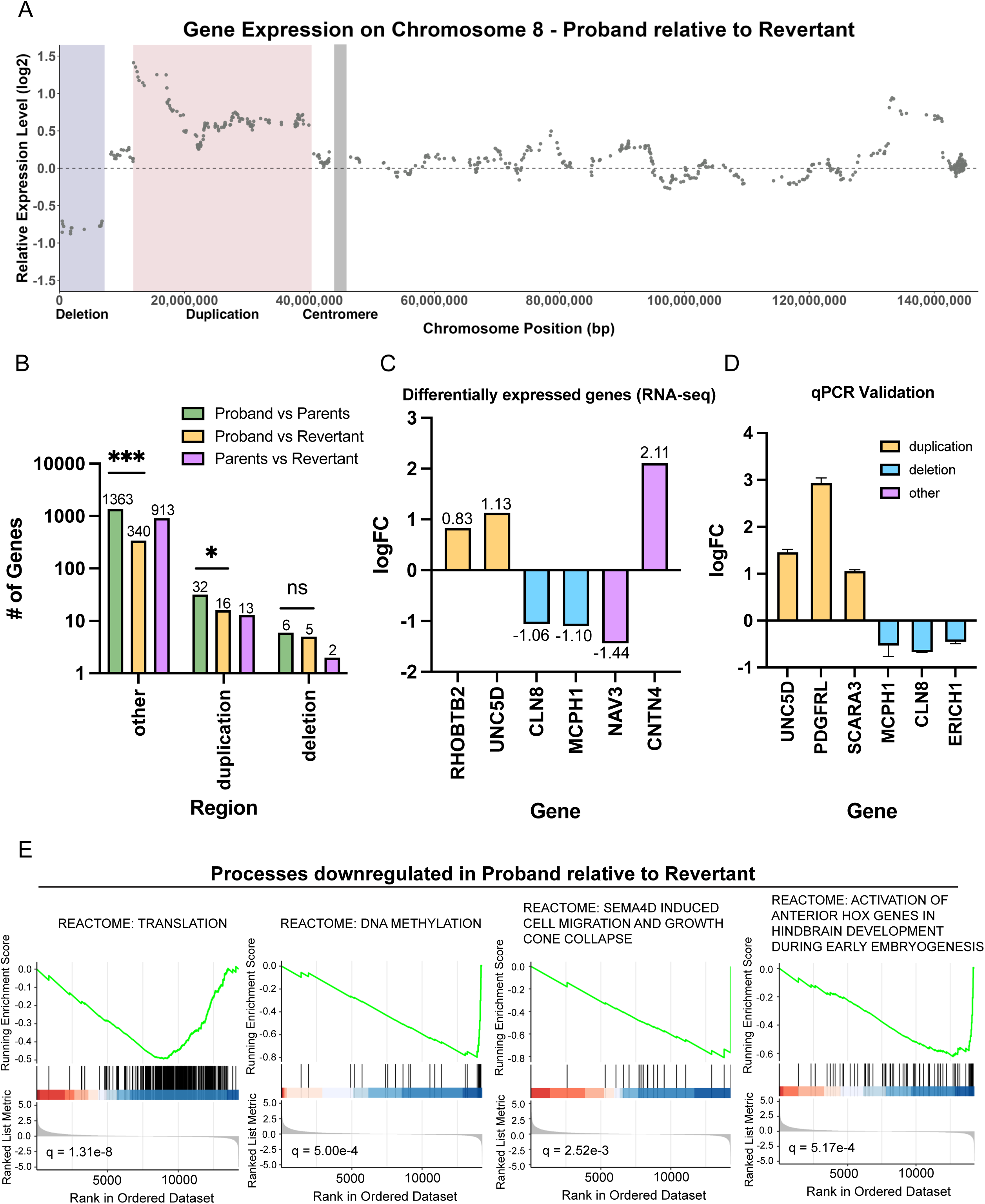
Comparing gene expression between proband and revertant iPS cells A. Diagram comparing relative gene expression vs chromosome position along Chromosome 8. The deleted region is highlighted in blue, the duplicated region is highlighted in red, and the position of the centromere is highlighted in gray. B. Bar graph displaying the number of differentially expressed genes according to the chromosomal region. Statistical testing was performed via Chi-squared test. ns, p ≥ 0.05; *, p ≤ 0.05; **, p ≤ 0.01; ***, p ≤ 0.001. C. Representative genes that were identified as differentially expressed between the proband and revertant iPS lines. A complete list of differentially expressed genes is included in Table S1. D. RT-qPCR validation of differentially expressed genes between proband and revertant iPS lines. Mean ± SEM, data from representative trials are shown (n ≥ 3 total trials). E. Representative biological pathways enriched among genes that were downregulated in the proband relative to the revertant. A complete list of GSEA results is included in Table S2.

We hypothesized that the isolation of a genetically-matched euploid revertant would allow us to precisely identify genes and processes dysregulated by the 8p rearrangement. In contrast, comparisons between the proband and the parents may be confounded both by genetic differences and by differences that arose during the cellular reprogramming process. We therefore performed genome-wide differential expression analysis between either the proband and the revertant or between the proband and the parents. Consistent with our hypothesis, we found that 1,401 genes were differentially expressed between the proband and the parents, and only 361 genes were differentially expressed between the proband and the revertant (Figure 4B). We speculate that these 361 genes are highly enriched for the genes that contribute to invdupdel(8p) pathology (Table S1). Notably, we also identified 928 differentially-expressed genes between the revertant and the parents – almost 3-fold more than we observed between the proband and the revertant – underscoring the significant impact that genetic confounding can have on the transcriptome (Figure 4B).

Within the deleted region, we observed that *CLN8* and *MCPH1* were significantly downregulated (Figure 4C). Mutations in *CLN8* have been reported to cause epilepsy and neurodegeneration (Sharkia et al. 2022). Mutations in *MCPH1* have been associated with microcephaly (Kristofova et al. 2022). Within the duplicated region, *RHOBTB2* and *UNC5D* were significantly overexpressed. *RHOBTB2* has been identified in prior studies as a potential candidate 8p driver gene due to its association with epileptic encephalopathy and other phenotypes overlapping with 8p syndrome (Okur et al. 2021). *UNC5D* overexpression has been associated with delayed migration of radial cortical neurons (Seiradake et al. 2014). qPCR of *CLN8*, *UNC5D*, and four other genes encoded within the rearranged region on 8p confirmed the changes in expression expected based on the copy number of these loci (Figure 4D).

Interestingly, we observed that a majority of differentially expressed genes were located outside of Chromosome 8p (Figure 4B). Notable genes included *NAV3*, which was downregulated, and *CNTN4*, which was upregulated (Figure 4C). Prior studies have linked *NAV3* to processes like neuronal cell migration; impairment in this process can contribute to agenesis of the corpus callosum and hypotonia (Ghaffar et al. 2024). *CNTN4* encodes a neuronal membrane protein that has roles in the plasticity and maintenance of neuronal networks. Duplications in *CNTN4* have been observed in cohorts of individuals with autism spectrum disorder (Glessner et al. 2009; Zhang et al. 2020).

Finally, we conducted gene set enrichment analysis (GSEA), comparing the proband and the revertant. We observed that certain fundamental biological processes, including protein translation and DNA methylation, were downregulated in the proband (Figure 4E and Table S2). Additionally, pathways related to neurodevelopment, including neural cell migration and hindbrain development, were enriched among downregulated genes. Fewer pathways were upregulated in the proband; the most significantly-upregulated process was homophilic cell adhesion (Table S2). We also repeated this analysis while excluding all genes encoded within the rearranged region of Chromosome 8p. We found a very strong overlap between the gene set analyses conducted with and without 8p; out of 187 gene sets significantly dysregulated in our initial GSEA, 176 of these gene sets remained significant at a Q < .05 threshold when 8p was excluded (Figure S3D-E and Table S3). These results are consistent with our finding that invdupdel(8p) results in the dysregulated expression of many more genes outside of 8p than on 8p (Figure 4B) and suggest that the *trans* effects of invdupdel(8p) are a significant contributor to the pathology of this condition.

### Transcriptional similarities between invdupdel(8p) and Trisomy 21

Previous reports have noted similarities between the clinical presentation of Down syndrome and other conditions caused by whole-chromosome aneuploidy (Krivega et al. 2022). On a cellular level, different autosomal trisomies result in similar changes in gene expression, which has been hypothesized to result from a shared stress response to aneuploidy (Dürrbaum et al. 2014; Sheltzer et al. 2012; Volk et al. 2017). Whether a complex chromosome rearrangement like invdupdel(8p) affects the transcriptome in a manner similar to a simple trisomy like Down syndrome is not known. To address this question, and to further explore the transcriptional consequences of invdupdel(8p), we compared the gene expression profile of the invdupdel(8p) proband, normalized to the revertant, to an iPS cell line with Trisomy 21, normalized to an isogenic diploid clone that arose spontaneously during passaging (Figure 5A) (MacLean et al. 2012; Meharena et al. 2022). As expected, we found that genes encoded on Chromosome 21 were expressed on average 54% higher in the Trisomy 21 line, while there was no consistent upregulation of Chromosome 21 genes in the invdupdel(8p) line (Figure 5B). To eliminate the *cis* effects resulting from aneuploidy, we excluded all genes encoded on Chromosome 21 and on the rearranged region of Chromosome 8p from our subsequent analyses.

**Figure 5:**
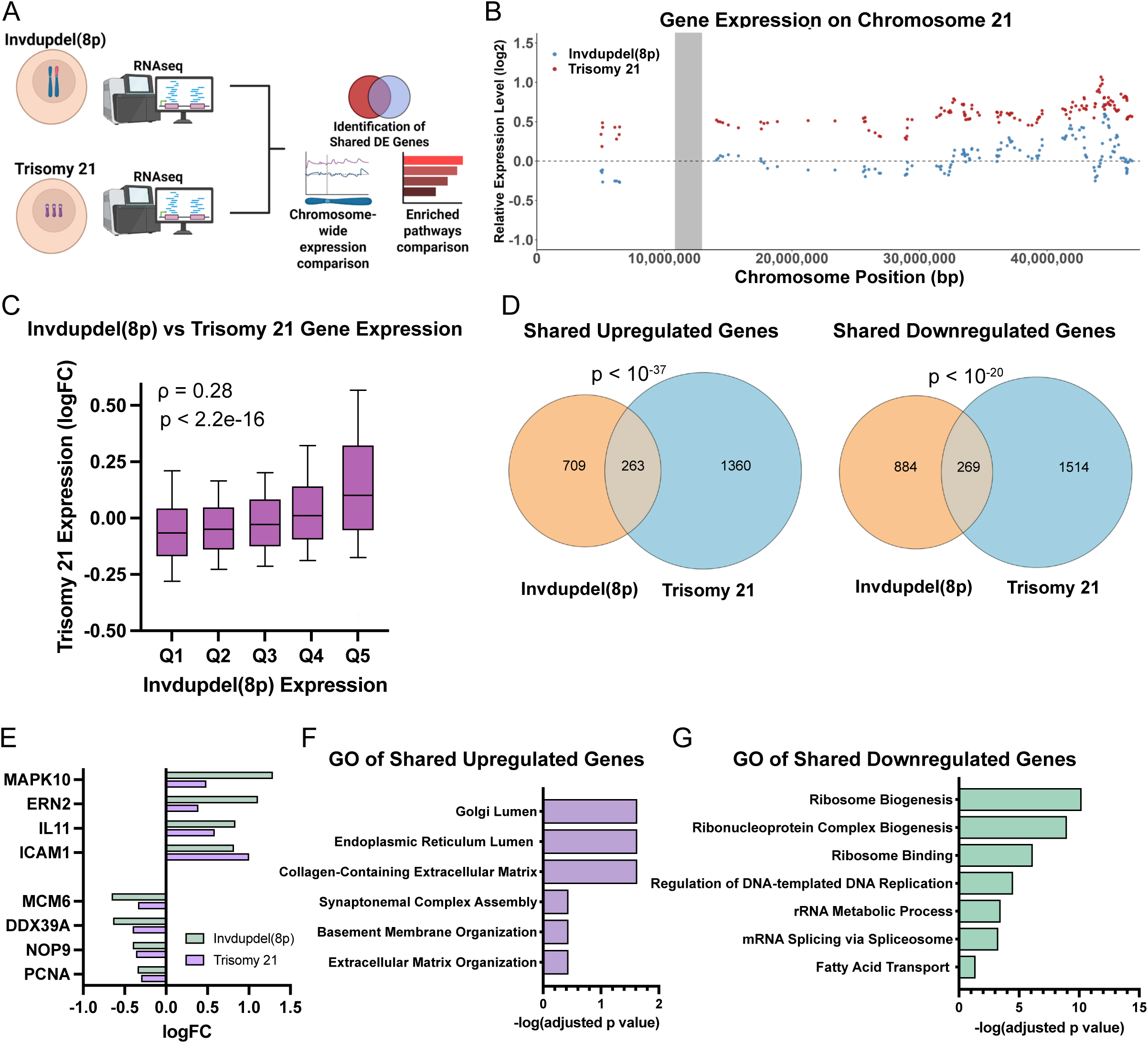
Identification of shared transcriptional changes between invdupdel(8p) and Trisomy 21 A. Schematic summarizing the comparative analysis and characterization of gene expression data from invdupdel(8p) and Trisomy 21 RNA-seq datasets. B. Diagram comparing relative gene expression vs chromosome position along Chromosome 21. C. X-axis displays quantile grouping of genes based on logFC of invdupdel(8p) analysis. Y-axis displays the quantile grouping of genes based on the logFC of the Trisomy 21 analysis. Spearman’s correlation coefficient and p value are displayed. D. Venn diagrams delineating shared upregulated (Left) and shared downregulated (right) DE genes between invdupdel(8p) and Trisomy 21 datasets. A complete list of genes is included in Table S4. E. Example genes that are differentially-expressed in both invdupdel(8p) and Trisomy 21 iPS cells. F. Gene Ontology of shared upregulated genes between invdupdel(8p) and Trisomy 21 datasets. Complete results are included in Table S5. G. Gene Ontology of shared downregulated genes between invdupdel(8p) and Trisomy 21 datasets. Complete results are included in Table S6.

Next, we sought to determine whether Trisomy 21 and invdupdel(8p) resulted in similar transcriptional changes. We found a highly-significant correlation in gene expression genome-wide between iPS cells derived from these two aneuploidy syndromes (rho = 0.28, P < 10^-15^; Figure 5C). 263 genes were upregulated and 269 genes were downregulated by both Trisomy 21 and invdupdel(8p), which are significantly greater than the overlaps expected by chance (P < 10^-37^ and P < 10^-20^, respectively, hypergeometric test; Figure 5D and Table S4). Transcripts upregulated by both conditions included the endoplasmic reticulum stress sensor *ERN2* and the apoptosis mediator *MAPK10*, while transcripts downregulated by both conditions included genes associated with cell proliferation (*PCNA*, *MCM6*), ribosome biogenesis (*NOP9*), and translation (*DDX39A*)(Figure 5E). Gene Ontology (GO) analysis revealed a significant upregulation of transcripts associated with the endoplasmic reticulum and the extracellular matrix, consistent with the induction of aneuploidy-induced protein folding stress (Donnelly and Storchová 2015). Downregulated GO terms were associated with translation, ribosome biogenesis, and the cell cycle, indicative of a general decrease in cell growth and metabolic activity (Figure 5F-G and Table S5-6). We also noted some GO terms that were specific to individual aneuploidies. For instance, we found that genes associated with the response to interferon were enriched among transcripts upregulated by Trisomy 21 and not invdupdel(8p), which is consistent with emerging data linking Down syndrome with dysregulated interferon signaling (Sullivan et al. 2016).

### Invdupdel(8p) antagonizes neural differentiation

As the revertant line can serve as an isogenic euploid control to the proband line, our chromosome engineering approach enables us to begin investigating how invdupdel(8p) alters neurodevelopment and causes other physiological changes associated with 8p syndrome. Toward this end, we performed neural differentiation (ND) on proband and revertant iPS cells, and we quantified the percentage of resulting neurons (MAP2^+^) and astroglia (S100B^+^) using automated image analysis with CellProfiler (Figure 6A-B) (Lickfett et al. 2022; McQuin et al. 2018). We confirmed that cell counts generated by CellProfiler correlated well with manual counts for nuclei (R=0.94), MAP2^+^ cells (R=0.81), and S100B^+^ cells (R=0.62)(Figure S4A-C).

**Figure 6:**
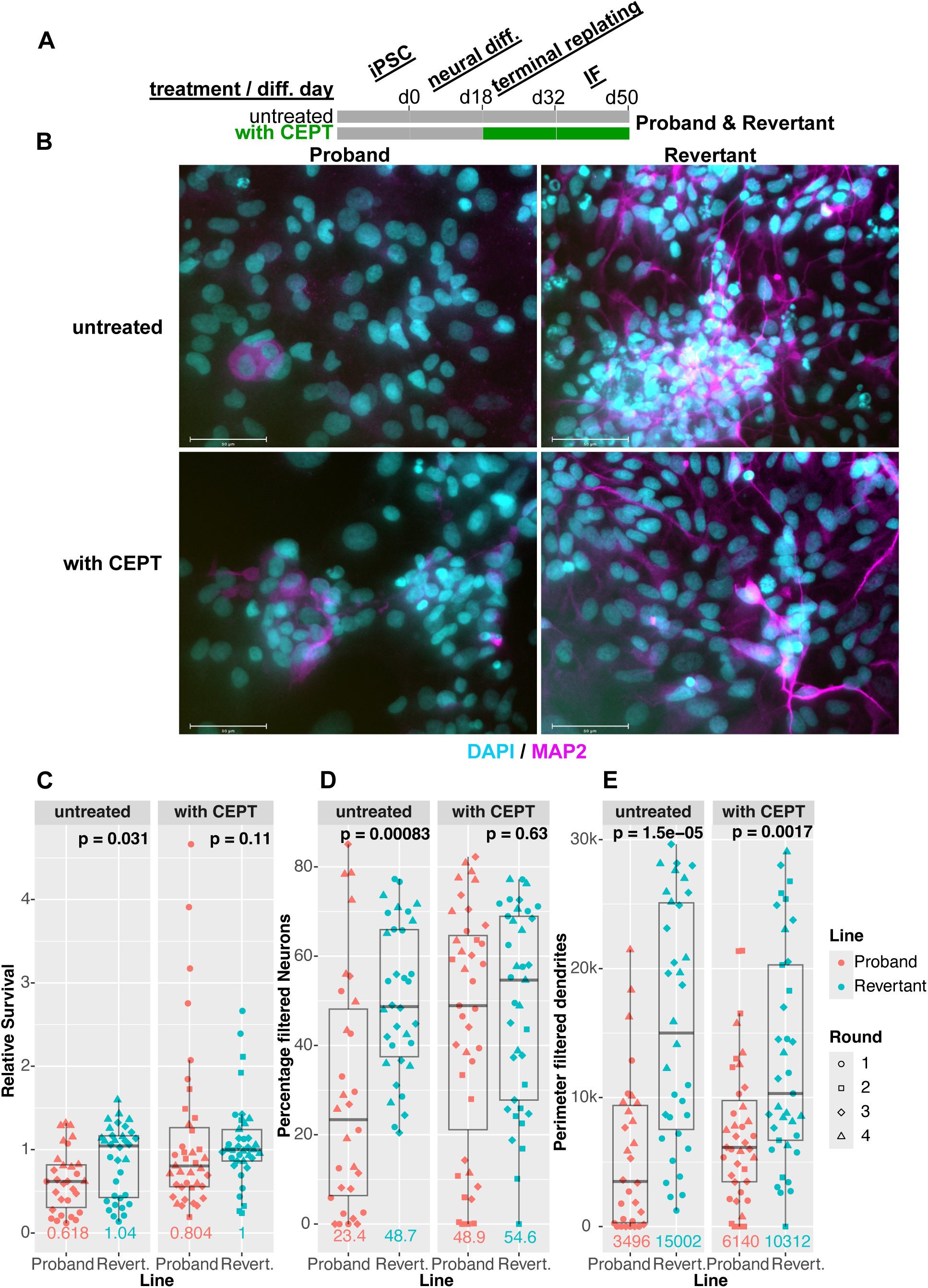
Analysis of neural differentiation in proband and revertant iPS cells A. Schematic of neural differentiation: proband and revertant iPS cells were differentiated as described in (Banda and Grabel 2014). On day 0 (d0) of neural differentiation, iPS media is replaced with N2B27 containing noggin (through d10). On d18, neural stem cells are terminally replated onto laminin-coated substrates and maintained in neural maturation media to the endpoint (d50). Following terminal replating, proband and revertant lines were cultured with (green) or without (gray) CEPT. B. Representative immunofluorescence images for neuronal marker MAP2 (magenta), and nuclei (cyan) counterstained with Hoechst-33342 (100 mm scale bars). Proband (left) and revertant (right) neural cultures differ in MAP2^+^ prevalence and morphology, with the latter showing extensive outgrowths indicative of neuronal maturation. CEPT-treated neural cultures (bottom) show increased cell numbers and MAP2^+^ prevalence in the proband line relative to untreated proband cultures (top). C. Relative cell survival was calculated by dividing the number of nuclei (range 40-700) per image (n = 9-14) by the median number of nuclei of the CEPT-treated revertant line in the corresponding round of differentiation. Colors denote proband and revertant lines, symbols distinguish four independent rounds of differentiation. *P-*values (two-tailed Wilcoxon rank-sum test) above revertant boxplots show the significance of median differences to the proband line, and medians are indicated below boxplots in their corresponding line color. D. The percentage of filtered neurons was calculated as the fraction of MAP2^+^ nuclei over the sum of all nuclei in each image and multiplied by 100. Colors, symbols, *P*-value, and median as indicated as in Fig 6C. E. The perimeter of filtered dendrites is the total length of the perimeter of the MAP2^+^ cells filtered as in (Lickfett et al. 2022) and described in the methods section. Colors, symbols, *P*-value and median as indicated as in Fig 6C.

We discovered that proband cultures exhibited a survival defect relative to the euploid revertant upon terminal re-plating during ND (P = 0.03, two-tailed Wilcoxon test; Figure 6C). Interestingly, addition of the cytoprotective small-molecule drug cocktail CEPT, which blocks apoptosis and inhibits stress-response signaling, rescued the viability difference between the proband and revertant cultures (Chen et al. 2021). Likewise, the percentage of MAP2^+^ neurons was significantly lower in proband relative to revertant cultures (P < 0.001; two-tailed Wilcoxon test), but this difference in neural differentiation was also suppressed by treatment with CEPT (Figure 6D). Correspondingly, at the end of the ND assay, many more proband cells remained unlabeled with either MAP2 or S100B compared to the revertant in untreated cultures but not in cultures exposed to CEPT (76% vs. 51%, P < .001, two-tailed Wilcoxon test; Figure S4D-E). While CEPT rescued the overall defects in cell survival and neural specification, we observed that MAP2 dendrites in the revertant line appeared consistently more mature and elongated relative to proband dendrites, even when accounting for the number of nuclei (Figure 6E and S4F). In total, these results reveal that the 8p rearrangement inhibits neural differentiation in iPS cells, but this defect can be partially rescued by a drug cocktail that enhances cell survival.

## Discussion

Here, we report the first successful isolation of a euploid clone derived from an individual with a complex chromosomal rearrangement. We achieved this through a two-step process, in which we established an intermediate cell population harboring a trisomy of the affected chromosome and then induced the loss of a single copy of the rearranged homologue. Prior work has demonstrated that prolonged passaging can be sufficient to restore euploidy in trisomic iPS cells; however, we did not observe spontaneous revertants from our invdupdel(8p) starting population. Whole chromosome aneuploidies induce a significant fitness cost in primary cells (Williams et al. 2008) and we speculate that the effects of invdupdel(8p) on iPS cell proliferation were not sufficient to allow the isolation of a spontaneous revertant population.

Instead, we used transient treatment with an TTK inhibitor to isolate a clone harboring a trisomy of Chromosome 8, and then we applied KaryoCreate to promote the missegregation of Chromosome 8 and facilitate the isolation of a euploid revertant (Bosco et al. 2023). KaryoCreate is one of several new strategies for targeted chromosome engineering that have recently been developed. Other methods include ReDACT, in which CRISPR is used to introduce a positive-negative selection cassette onto an aneuploid chromosome, and a kinesin-based approach in which a plant kinesin is used to drag a chromosome to the wrong centrosome (Girish et al. 2023; Tovini et al. 2023; Truong et al. 2023). We elected to use KaryoCreate, as we anticipated that it would produce less DNA damage than these other methods and because it does not require stable modification of the target cell’s genome. Our work verifies the earlier publication from the Davoli Lab and suggests that KaryoCreate can be an effective strategy for inducing chromosome-specific missegregation events.

Gene expression analysis performed across the proband, parental, and revertant iPS cells revealed many fewer differentially-expressed genes when the proband and the revertant were directly compared. We speculate that this is because the proband and the revertant are isogenic while the proband and each parent share only 50% genetic similarity. Our expression analysis revealed 361 genes that were differentially expressed between the proband and the revertant. Notably, more than 300 of these genes were located outside the invdupdel(8p) region, highlighting how this chromosomal rearrangement can impact the transcriptional landscape throughout the genome. We recognize that, even in an isogenic background, some of these gene expression changes could occur due to the single-cell cloning process or as a consequence of mis-segregating Chromosome 8. However, our comparison of the invdupdel(8p) transcriptome with that of Trisomy 21 reveals a striking and significant convergence in the cellular response to these distinct aneuploidies. Despite involving different chromosomes and rearrangement complexities, both conditions trigger a common transcriptional signature characterized by the downregulation of genes involved in fundamental processes like translation and ribosome biogenesis, and the upregulation of genes related to the extracellular matrix. This shared response provides additional evidence for the existence of a general cellular stress mechanism activated by gene dosage imbalances, regardless of the specific genes involved in the primary aneuploidy (Dürrbaum et al. 2014; Sheltzer et al. 2012; Sheltzer 2013).

By taking advantage of our engineered revertant, we investigated how invdupdel(8p) impacts neural differentiation in a genetically-controlled context. In these assays, the proband exhibited a significant defect in both survival and neuronal (MAP2^+^) specification. Inclusion of the cytoprotective small-molecule cocktail CEPT (chroman-1, emricasan, polyamines, and trans-ISRIB) upon terminal plating of neural progenitors rescued both proband cell survival and neuronal differentiation (Chen et al. 2021). CEPT supports cell attachment, inhibits apoptosis, and reduces cell stress, which suggests a number of ways in which the proband cells may be more vulnerable than revertant cells during ND. To our knowledge, our work is the first to demonstrate that defective survival and differentiation in an aneuploid iPS line can be mitigated by treatment with CEPT. As our transcriptomic analysis highlighted similarities in the expression signatures of invdupdel(8p) and Trisomy 21, this shared molecular pathology suggests that CEPT may be a broadly useful tool to enhance ND in Down syndrome and other aneuploidy conditions. However, even with inclusion of CEPT, proband neurons extended fewer maturing, elongated dendrites than revertant cells. This indicates that the impact of the invdupdel (8p) rearrangement on proband iPSC-derived neurons may extend beyond simple cell stress during terminal replating and indicates some CEPT-resistant cellular phenotypes for follow-up studies.

Finally, chromosome engineering represents a powerful potential approach for treating multiple disorders caused by chromosomal rearrangements and copy number changes. Prior investigators have silenced or eliminated simple trisomies and chromosomal duplications in tissue culture through single genetic manipulations (Inoue et al. 2019; Li et al. 2012; Munezane et al. 2024),. To our knowledge, the work described in this manuscript represents the first instance in which a complex chromosome rearrangement has been genetically eliminated from human cells. Aneuploidy is associated with both genetic variability (e.g., in the size of the duplicated and deletion region on Chromosome 8p) (Okur et al. 2021) as well as non-genetic variability (e.g., due to stochastic differences in cell signaling that impact the robustness of biological pathways) (Beach et al. 2017). Chromosome engineering to restore euploidy could potentially represent a “one size fits all” approach to address syndromes caused by chromosomal alterations that encompass different regions and produce different molecular sequelae.

However, we were only able to produce a euploid revertant via a two-step process that included a trisomic intermediate. The currently-laborious nature of our protocol represents an important challenge for successful clinical translation. Furthermore, we note that our euploid revertant now harbors an isodisomy of Chromosome 8. While this approach avoids the introduction of foreign genetic material that could potentially produce immunogenic neoepitopes, epigenetic imprinting on a targeted chromosome could result in unintended functional consequences. Isodisomy of Chromosome 8 is compatible with normal development (Benlian et al. 1996; Yu et al. 2022), though this tolerability may not extend to other chromosomes that are subject to significant imprinting.

In future work, the establishment of improved techniques for chromosome engineering, coupled with a comprehensive analysis of the effects of these manipulations on chromosomal imprinting, could facilitate the development of new therapeutic strategies to treat 8p syndrome and other aneuploidy-associated disorders with chromosome therapy.

## Methods

### Acquisition of biological specimens

iPS cells from an individual with invdupdel(8p) as well as two parental iPS cell lines were provided by Project 8p (Shah 2024). Cell lines were received at passage 13. STR profiling was performed at the University of Arizona Genetics Core facility and verified the relationship between parental and proband cell lines. Research was conducted in accordance with Yale University and NIH guidelines (IRB Protocol ID: 2000033853).

### Cell culture conditions

iPS cells were cultured in mTeSR Plus (STEMCELL Technologies, cat. no. 100-0276), on plates coated in Matrigel (Corning, cat. no. 356231). Routine colony passaging and dissociation were done using Dispase (STEMCELL Technologies, cat. no. 07913). Media was changed daily, and cells were passaged every 4 days. For single cell applications, cells were dissociated using Accutase (STEMCELL Technologies, cat. no. 07920) and plated on Matrigel coated plates containing mTeSR Plus supplemented with 10 µM Y-27632 (STEMCELL Technologies, cat. no. 72304). To assess cell proliferation, growth curves, and doubling times in vitro, 150,000 cells were seeded per well in 4 wells of a 12-well plate. After 24 hours, the media was refreshed, and cells were transferred to the Sartorius Incucyte S3 Live-Cell Analysis System for real-time imaging. Cells were imaged over a 56-hour period (4 or 3 wells per cell type, 16 images per well), with media changes performed every other day. Cell confluence over time was quantified using Incucyte software and used to generate growth curves. Doubling times were calculated for each image, and only growth curves with R² ≥ 0.97.5 were included in the analysis. Cells were maintained at 5% CO_2_, 5% O_2_, 37° C in a humidified chamber.

### TaqMan copy number analysis

Genomic DNA from iPS cells was extracted using Qiagen DNeasy Blood & Tissue Kit (cat. no. 69506). Each sample was loaded in quadruplicate in 384 well plates. Quantitative PCR was performed using TaqPath ProAmp Master Mix (Applied Biosystems, cat. no. A30867). A probe targeting *RPP40* (Applied Biosystems, cat. no. 4316831) was used as a reference assay. Other TaqMan copy number probes from Applied Biosystems that were used in this work include: *MICU3* (Assay ID: Hs06219542_cn, location: Chr.8:17109660), *CSMD1* (Assay ID: Hs06164679_cn, location: Chr.8:2944759), and *CRISPLD1* (Assay ID: Hs02746949_cn, location: Chr.8:75032222).

Genomic copy number was typically calculated by normalizing to MCF10A, a human mammary epithelial cell line with a near-diploid karyotype (Girish et al. 2023). For the analysis in Figure 3B, *CRISPLD1* was normalized to the proband rather than to MCF10A.

### Cloning of mCherry into KNL1^Mut^-dCas9 KaryoCreate Plasmid

To facilitate the isolation of iPS cells transiently transfected with the KaryoCreate system, we introduced a fluorescent vector into the KNL1^Mut^-dCas9 plasmid. The blasticidin resistance marker in pHAGE-KNL1^Mut^-dCas9 was replaced with the mCherry construct from LRCherry2.1 (Addgene #108099) via HiFi assembly (New England Biolabs, cat. no. E2621L), using PCR primers atagctaggtagcctagagggcccgcggttACTTTGGCGCCGGCT and atattcatttctttgcaagttaCTGAATAATAAGATGACATGAACTACTATACGTACTGC to generate the mCherry-KNL1^Mut^-dCas9 plasmid (Addgene: 229940). The plasmid sequence was verified using whole-plasmid sequencing (Plasmidsaurus).

### Generation of trisomic proband intermediate cells

We speculated that the rate of spontaneous chromosome missegregation in the proband was insufficient to allow the generation of revertant clones during spontaneous passaging. While we did not attempt to calculate an exact chromosome missegregation rate in the proband, we elected to use the TTK inhibitor AZ3146 (Selleck Chemicals, cat. no. S2731) to induce a transient period of elevated chromosomal instability. Based on prior experiments testing the effects of AZ3146 treatment, (Lukow et al. 2021; Vasudevan et al. 2020) we elected to use 1.5 µM AZ3146. 2 million cells were treated with AZ3146 for 48 hours, and then the drug was washed out. 24 hours after drug washout, the cells were harvested and single-cell sorted. 8p copy number was determined via qPCR in approximately 60 clones, and two clones that harbored a trisomy of Chromosome 8 were identified and retained for further characterization.

### Generation of a revertant iPS cell line

Approximately 1 x 10^6^ trisomic proband iPS cells were transiently co-transfected with a plasmid that expressed a gRNA (Addgene: 229941) specific to the Chromosome 8 centromere region and the mCherry-KNL1^Mut^-dCas9 plasmid. Proband cells were transfected using the Amaxa 4D-Nucleofector X (Lonza) system. After nucleofection, the buffer mixture containing cells was seeded onto two wells of a Matrigel-coated six-well dish. 8p copy number was determined via qPCR in approximately 100 clones, and one clone in which disomy of Chromosome 8 had been restored was retained for further characterization.

### RNA-seq

Total cellular RNA was isolated using the Qiagen RNeasy Mini Kit (Qiagen, cat. no. 74106) and submitted to Novogene for RNA-seq and quantification. Raw data in FASTQ format were processed using Novogene’s analysis pipeline to generate raw read counts. Read counts were filtered to remove lowly expressed genes and normalized across samples using TMM normalization with edgeR (v4.2.1) (Chen et al. 2024). Differential expression analysis was performed with edgeR (v4.2.1). Samples extracted from the mother’s iPS cells and the father’s iPS cells were combined into a group labeled “Parents”. For the analysis in Figure 4B, an additional threshold of absolute log_2_ fold change > 1 was applied to identify differentially expressed (DE) genes.

CPM counts were calculated and transformed to a log_2_ scale using edgeR (v4.2.1) and ranked using the Signal2Noise metric with the local version of the GSEA tool (v4.3.3) http://www.broadinstitute.org/gsea/index.jsp. Gene Set Enrichment Analysis (GSEA) was performed using the Hallmark, C2, and C5 gene sets from MSigDB, with clusterProfiler (v4.12.6). Statistical testing was performed using a two-tailed Pearson’s Chi-Square test. Analyses were performed using R Statistical Software (v4.4.1; R Core Team 2024) and tidyverse (v2.0.0) (R Core Team 2024; Wickham et al. 2019, 2023b, 2023a, 2024). Gene expression data generated in this study is available at GSE281425.

### Quantitative real-time qPCR

Total cellular RNA was extracted using the Qiagen RNeasy Mini Kit (Qiagen, cat. no. 74106). cDNA was synthesized using SuperScriptIV VILO Master Mix (Invitrogen, cat. no. 11756500). Quantitiative PCR was performed for selected targets using PowerTrack SYBR Green Master Mix (Applied Biosystems, cat. no. A46109) and quantified using the QuantStudio 6 Flex Real-Time PCR system (Applied Biosystems). qPCR primers are listed in Table S7.

### Comparison of gene expression data from invdupdel(8p) and Trisomy 21

Raw RNA-seq data (FASTQ) for the Trisomy 21 iPS cell dataset (GSE185192) were retrieved from the Sequence Read Archive (Katz et al. 2022). Quality control was performed using FastQC (v0.12.1) (Babraham Bioinformatics 2023) and fastp (0.23.2) (Chen 2023), and reads were aligned to the human reference genome (GRCh38) using HISAT2(2.2.1) (Kim et al. 2019). Gene-level quantification was conducted with featureCounts (Subread 2.0.3) (Liao et al. 2014), following the workflow used in Novogene’s standard RNA-seq pipeline, to ensure compatibility with analyses of the invdupdel(8p) dataset. Differential gene expression analysis was performed using edgeR (v4.2.1) (Chen et al. 2024), following the same processing pipeline as used for the invdupdel(8p) dataset. Gene Ontology (GO) enrichment analyses were conducted for both the Trisomy 21 and invdupdel(8p) datasets using Enrichr (https://maayanlab.cloud/Enrichr/) (Xie et al. 2021), with condition-specific background sets and a significance threshold of FDR ≤ 0.05 for defining differentially expressed genes.

### Whole genome sequencing and copy number alteration detection

Genomic DNA was isolated as described above. Sequencing was performed using Short-Read Human WGS services provided by GENEWIZ from Azenta Life Sciences, as described (https://www.genewiz.com). After sequencing, copy number alterations in the samples were inferred by a custom Nextflow wrapper (v1.0) for CNVKit (v0.9.11) with “batch--method wgs” command, specifying a bin size of 100,000 (Olshen et al. 2011; Talevich et al. 2016; Venkatraman and Olshen 2007). For selected samples, a bin size of 50,000 was used to improve segmentation sensitivity and copy number estimation. Absolute copy numbers of segments were inferred by CNVkit with “call” command, while copy numbers of the bins were derived from log_2_ copy ratios. Plots of copy number alterations were generated using a custom Python script (v3.12.4) with Matplotlib (v3.8.4) and pandas (v2.2.2) (Hunter 2007; The pandas Development Team 2024; The Python Software Foundation 2024). The script was adapted from CNVkit’s scatter.py with specific modifications to enhance visual representation. This low-depth analysis was designed to be an economical strategy to identify somatic copy-number alterations throughout the genome, but we recognize that that small copy-number alterations or other complex rearrangements may not be detected at this resolution. The full details of the modifications, as well as the complete workflow, are available in a public repository on GitHub https://github.com/sheltzer-lab/Karyotyping.

### Neural differentiation

Proband and revertant iPSC lines were differentiated using a monolayer neural differentiation method (Banda and Grabel 2014). iPSCs were cultured on mouse embryonic feeder (mEF) layers in iPSC media (DMEM/F-12 (Life Technologies, cat. no. 11330-032), KOSR (Life Technologies, cat. no. 10828-028), bFGF (final concentration 4ng/mL; Life Technologies, cat. no. PHG0264), Glutamax (Life Technologies, cat. no. 35050-061), NEAA (Life Technologies, cat. no. 11140-050) and ß-mercaptoethanol (Life Technologies, cat. no. 21985-023). On day 3 following passage, media was changed from iPSC media to N2B27: Neurobasal media (Life Technologies, cat. no. 21103-049), N2 (25 μg/ml insulin, 100 μg/ml holo-transferrin, 100 μM putrescine, 20 nM progesterone, and 30 nM selenite), B27 (Life Technologies, cat. no. 12587-010), Glutamax (Life Technologies, cat. no. 35050-061), Penicillin-Streptomycin (151040-122, cat. no. Life Technologies). Media was changed every other day and supplemented with noggin (R&D Systems, cat. no. 3344-NG-050) through day 10 (D10). After D10, media with N2B27 was changed every day until D18. At D18, cultures were enzymatically passaged with Accutase (EMD Millipore, cat. no. SCR005) onto poly-D lysine and laminin-coated substrates at 2.1 x 10^4^ cells/cm^2^ into Neural Differentiation Media (NDM): Neurobasal Media, B27, Glutamax, NEAA, Penicillin-Streptomycin supplemented with brain-derived neurotrophic factor (BDNF; 10 ng/ml; Peprotech), glial-derived neurotrophic factor (10 ng/ml; Peprotech), neurotrophin-3 (10ng/ml; Peprotech). For cultures receiving CEPT (Tocris, cat. no. 7991), CEPT was added from D18, following cell replating.

### Immunofluorescence

Neural differentiation cultures were fixed in 4% paraformaldehyde (Electron Microscopy Sciences) at room temperature (RT) between D35 and D50 of differentiation. Following fixation, slides were rinsed in PBS three times for 5 minutes each, permeabilized in 1% Triton-X in PBS for 10 minutes at RT and then blocked for 1 hour at RT in blocking buffer containing 0.1% Triton-X, 5% bovine serum albumin, and 2% normal goat serum. Primary antibodies for MAP2 (Abcam; ab5392; 1:10,000) and S100B (Sigma-Aldrich; s2532; 1:1000) were diluted and hybridized in blocking buffer overnight at 4°C. Slides were washed with 0.1% Triton-X in PBS (wash buffer) 3 times for 5 minutes each at RT. Secondary antibody labeling was performed by diluting Alexa Fluor secondaries (Life Technologies) at 1:500 in blocking buffer for 1 hour at RT. Slides were washed three times for 5 minutes each in wash buffer before counterstaining nuclei with Hoechst 33342 (1:10,000 for 10 minutes). Cells were mounted in Prolong Antifade Mountant (ThermoFisher Scientific) and imaged on the EVOS FL Auto system (ThermoFisher Scientific).

### Image analysis

For each round of neural differentiation (ND), cell line (proband and revertant iPSC), and condition (untreated vs. CEPT), images were quantified by automated image analysis using CellProfiler (v4.2.4) (McQuin et al. 2018). Nuclei were identified using the IdentifyPrimaryObjects module (Otsu 3-class, Hoechst minimum of 0.0024, diameter range: 20-50, threshold smoothing scale: 1.3488, and size of smoothing filter 6). Nuclei outside the diameter range or touching image borders were discarded. Primary object counts were validated by comparison to manual nuclei counts in ND5/6 (Pearson R ≥ 0.9). Nuclei were expanded by 3 pixels (ExpandOrShrinkObjects) and masked (MaskObjects) by minimal (fraction 0.25) overlap with round-specific, log-transformed and thresholded MAP2 (ND1/2: 0.0024, ND3/4: 0.003) or S100B (ND1/2: 0.0024, ND3/4: 0.0036).

IdentifySecondaryObjects, IdentifyTertiaryObjects and SplitorMergeObjects were used to identify cell soma and map dendrites, as adapted from (Lickfett et al. 2022). Cells were filtered for MAP2 and S100B minimum thresholds above (FilterObjects), and thereby classified as either unlabeled, neuron (MAP2^+^) or astrocyte (S100B^+^). Neuronal soma were filtered with a minimal eccentricity (0.8, the ratio of the distance between the foci of the ellipse and its major axis length) and a maximal (5 pixel) median radius of any pixel within to the closest pixel outside, to identify elongated dendrites, as in (Lickfett et al. 2022).

### Data Visualization

Scientific illustrations were assembled using BioRender. Graphs were generated using GraphPad Prism. GSEA plots were generated using enrichplot (v1.24.4) (Yu 2024). Plots were created using ggplot2 (v3.5.1), with ggthemes (v5.1.0), MetBrewer (v0.2.0) and eularr (v7.0.2) used as well, (Arnold et al. 2024; Mills 2022; Wickham 2016).

## Data access

All raw and processed RNA-seq data generated in this study have been submitted to the NCBI Gene Expression Omnibus (GSE281425). All code for data analysis (including supplemental code) is available at github.com/sheltzer-lab/8p-rearrangement.

## Acknowledgements

We are grateful to Project 8p and to the anonymous donors who provided the biological specimens used in this analysis. This work was supported in part by a research grant from Project 8p. Research in the Sheltzer Lab is supported by NIH grants R01CA237652 and R01CA276666, Department of Defense grant W81XWH-20-1-068, an American Cancer Society Research Scholar Grant, and a sponsored research agreement from Ono Pharmaceuticals. Research in the Pinter Lab is supported by NIH grant R35GM153326. Research in the Davoli Lab is supported by NIH grant NIH R01DK135089. We thank members of the Yale hESC/iPSC Core Facility for providing reagents and assistance with iPS cell culture. Additionally, we thank the Yale Flow Cytometry Core for their assistance with cell sorting. The Core is supported in part by an NCI Cancer Center Support Grant # NIH P30 CA016359. The BD Symphony was funded by shared instrument grant # NIH S10 OD026996. Work in the Davoli Lab is supported by NIH grant R37CA248631.

## Author Contributions

S.N.L. performed the chromosome engineering experiments and iPS cell manipulations, analyzed the resulting data, and contributed to writing and editing the manuscript. E.C.B. performed the neural differentiations, immunofluorescence imaging, and manual quantification. L.Q. analyzed the whole-genome sequencing and RNA-seq data and compared the effects of invdupdel(8p) to Trisomy 21. S.L.T. assisted with the iPS cell passaging and the overall experimental approach. R.A.H. assisted with the analysis of the whole-genome sequencing data. T.D. provided reagents and assistance for the experiments involving KaryoCreate. E.C.B. and S.F.P. developed and tested the CellProfiler pipeline, generated figures and contributed to writing and editing the manuscript. J.M.S. designed the project, supervised the research, analyzed the data, and contributed to writing and editing the manuscript.

