## Supplemental figures for "Chromosome engineering to correct a complex rearrangement on Chromosome 8 reveals the effects of 8p syndrome on gene expression and neural differentiation"

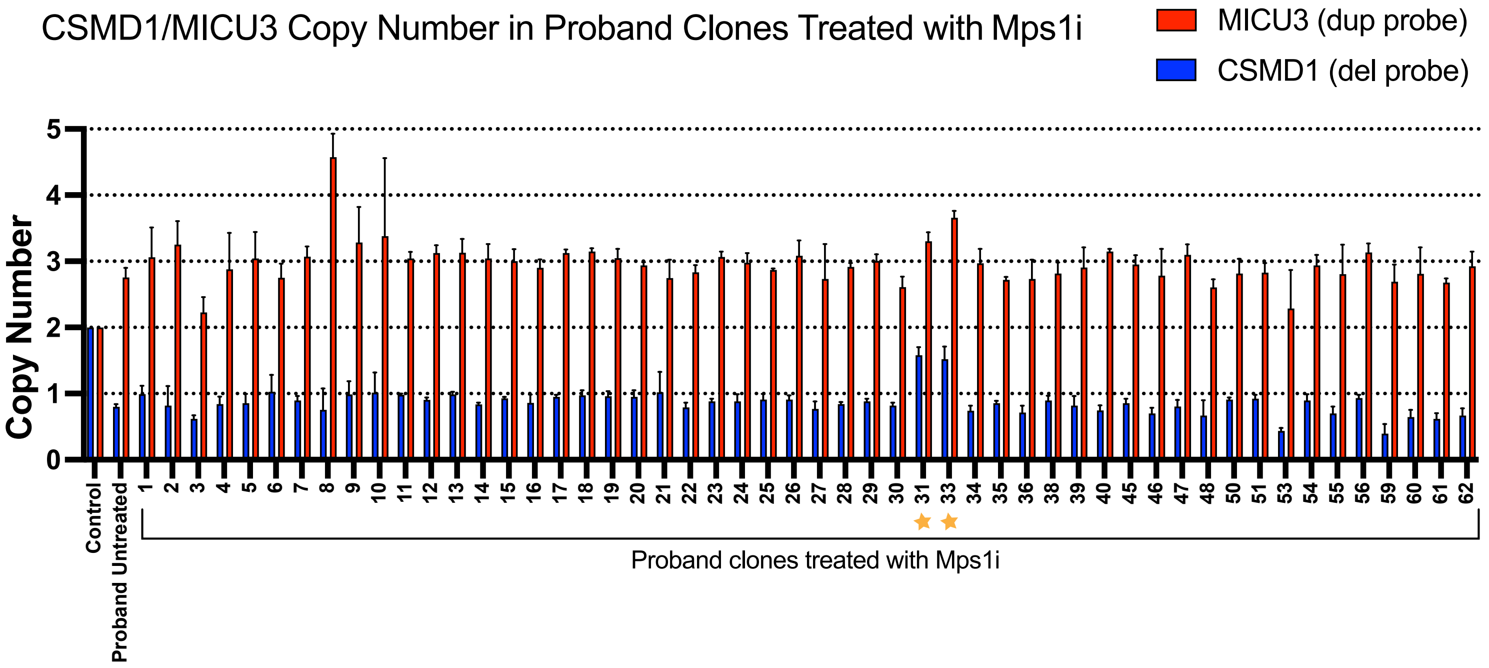


**Supp. Figure S1: Copy number analysis of proband clones after treatment with AZ3146**
Copy numbers of *CSMD1* and *MICU3* in clones treated with AZ3146. Trisomic clones 31 and 33 are labeled with stars. Mean ± SEM, data from representative trials are shown (n ≥ 3 total trials).

Figure S2

A

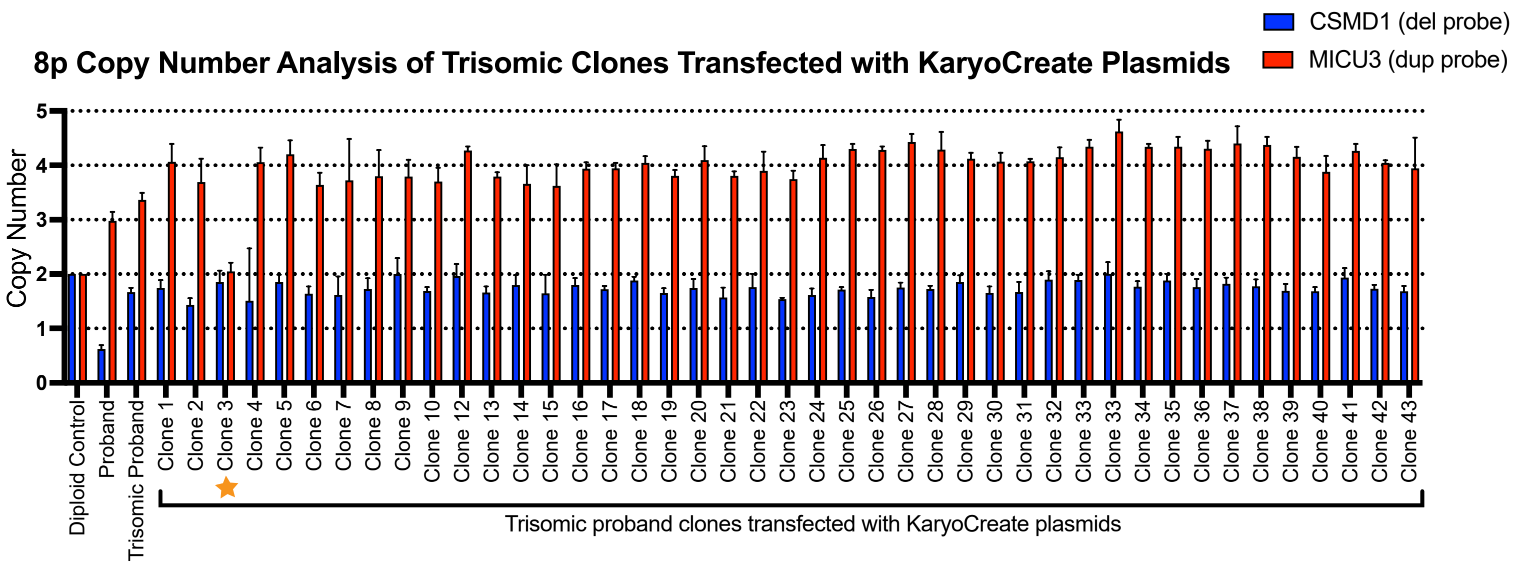


B

**STR Profiles**

| Locus | Parent 1 | | Parent 2 | | Proband | | Reverted Proband | |
|---|---|---|---|---|---|---|---|---|
| D5S818 | 12 | 12 | 11 | 11 | 11 | 12 | 11 | 12 |
| D13S317 | 9 | 12 | 12 | 12 | 9 | 12 | 9 | 12 |
| D7S820 | 10 | 10 | 8 | 9 | 9 | 10 | 9 | 10 |
| D16S539 | 11 | 13 | 13 | 13 | 13 | 13 | 13 | 13 |
| VWA | 17 | 18 | 16 | 17 | 17 | 17 | 17 | 17 |
| TH01 | 6 | 9.3 | 6 | 9 | 9 | 9.3 | 9 | 9.3 |
| AM | X | X | X | Y | X | X | X | X |
| TPOX | 8 | 11 | 11 | 11 | 11 | 11 | 11 | 11 |
| CSF1PO | 12 | 13 | 10 | 10 | 10 | 12 | 10 | 12 |
| D3S1358 | 15 | 15 | 17 | 19.1 | 15 | 19.1 | 15 | 19 |
| D21S11 | 29 | 33.2 | 28 | 29 | 28 | 29 | 28 | 29 |
| D18S51 | 14 | 14 | 13 | 15 | 13 | 14 | 13 | 14 |
| Penta_E | 5 | 15 | 7 | 13 | 7 | 15 | 7 | 15 |
| Penta_D | 9 | 9 | 12 | 14 | 9 | 12 | 9 | 12 |
| D8S1179 | 8 | 17 | 9 | 10 | 10 | 17 | 10 | 10 |
| FGA | 21 | 22 | 23 | 24 | 21 | 24 | 21 | 24 |

C

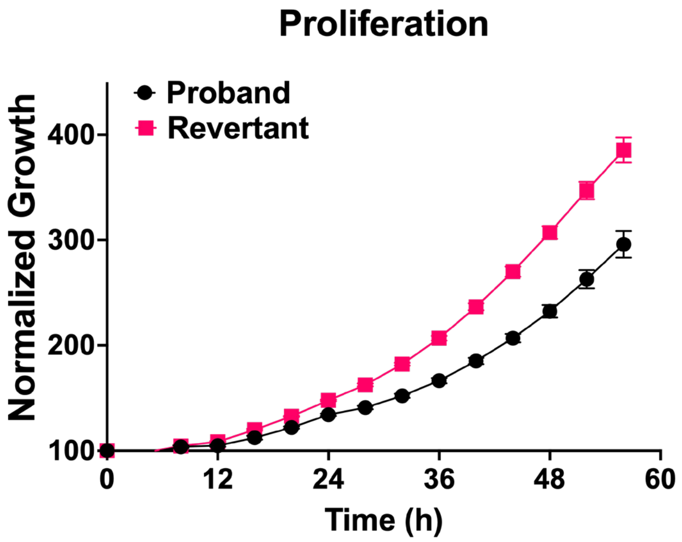


**Supp. Figure S2:** Analysis of trisomic clones transfected with KaryoCreate plasmids
**(A)** Copy numbers of *CSMD1* and *MICU3* in trisomic clones after transient transfection with KaryoCreate plasmids. Revertant clone 31.3 is labeled with a star. Mean ± SEM, data from representative trials are shown (n ≥ 3 total trials). Genomic DNA extracted from MCF10A cells was used as a diploid control.
**(B)** STR analysis of the parental, proband, and revertant cells. The D8S1179 locus, which is located on Chromosome 8, is highlighted in yellow.
**(C)** Growth curves of revertant clone 31.3 and proband iPS cell lines.

Figure S3

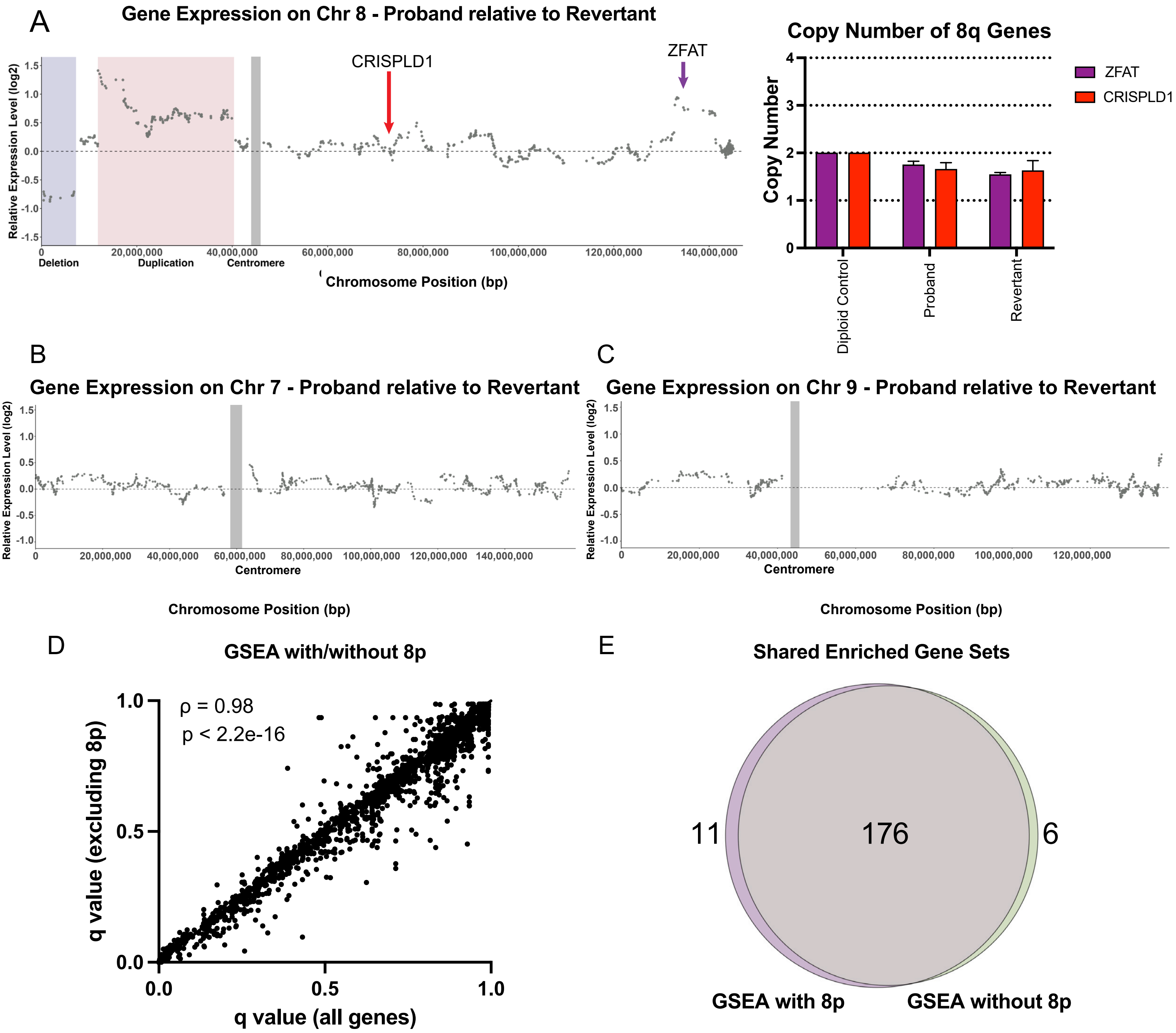


**Supp. Figure S3: Additional analysis of gene expression in invdupdel(8p) cells**

**(A)** Left: Diagram comparing relative gene expression vs chromosome position along Chromosome 8. The deleted region is highlighted in blue, the duplicated region is highlighted in red, and the position of the centromere is highlighted in gray. Red arrow delineates binding location of *CRISPLD1* probe. Purple arrow delineates binding location of *ZFAT* probe. Right: Copy numbers of *ZFAT* and *CRISPLD1*. Mean ± SEM, data from representative trials are shown (n ≥ 3 total trials).

**(B)** Diagram comparing relative gene expression vs chromosome position along Chromosome 7.

**(C)** Diagram comparing relative gene expression vs hromosome position along Chromosome 9.

Spearman's rank correlation between GSEA analyses either including or excluding genes contained within the invdup-del(8p) rearrangement. Complete results are included in Table S3.

**(D)** Venn diagram displaying shared enriched gene sets between GSEA including or excluding genes contained within the invdupdel(8p) rearrangement.

Figure S4

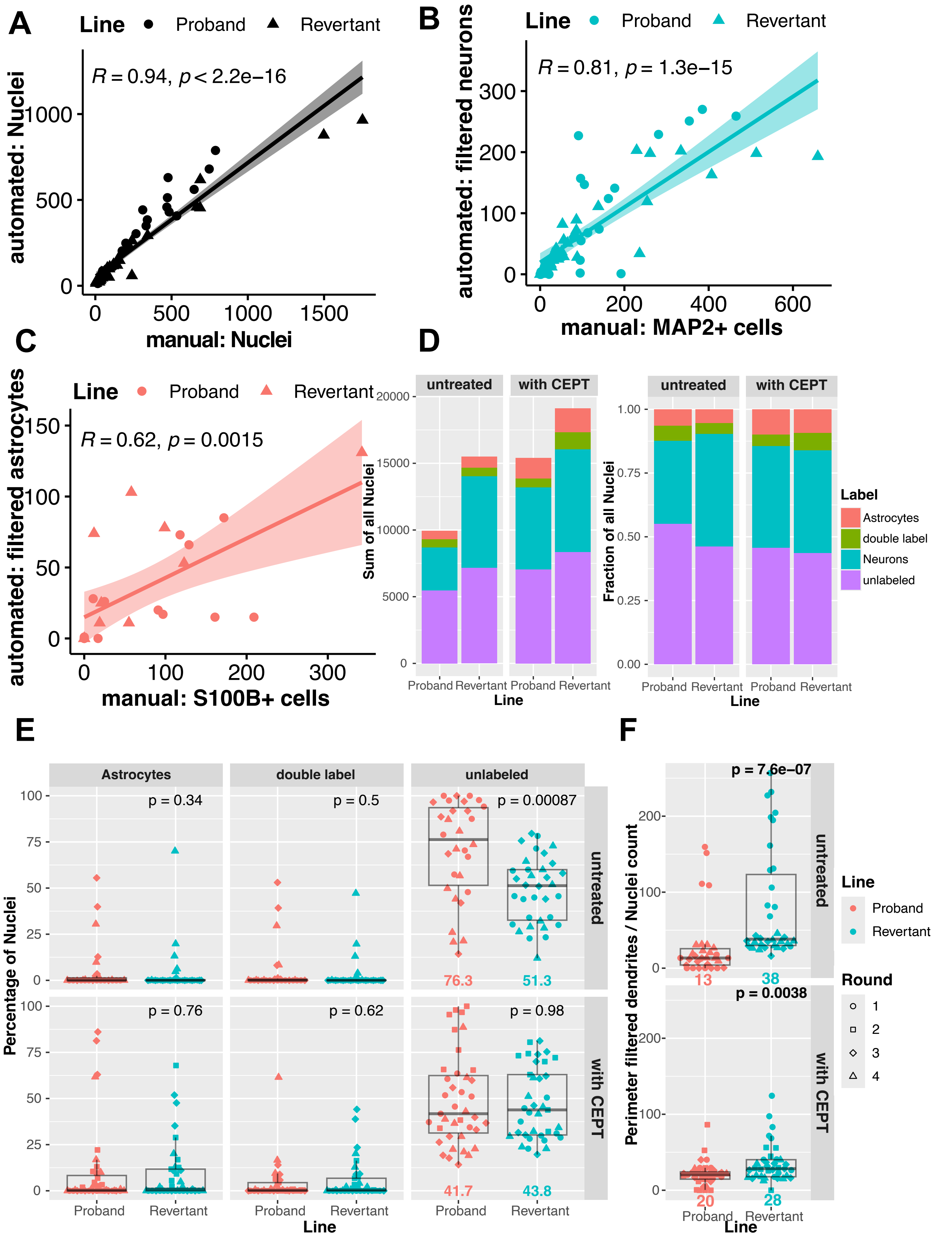

A
Line Proband Revertant
R = 0.94, p < 2.2e−16
automated: Nuclei
manual: Nuclei
B
Line Proband Revertant
R = 0.81, p = 1.3e−15
automated: filtered neurons
manual: MAP2+ cells
C
Line Proband Revertant
R = 0.62, p = 0.0015
automated: filtered astrocytes
manual: S100B+ cells
D
untreated
with CEPT
Sum of all Nuclei
Fraction of all Nuclei
Line
Label
Astrocytes
double label
Neurons
unlabeled
E
Astrocytes
double label
unlabeled
p = 0.34
p = 0.5
p = 0.00087
p = 0.76
p = 0.62
p = 0.98
76.3
51.3
41.7
43.8
Percentage of Nuclei
Proband
Revertant
Line
F
p = 7.6e−07
p = 0.0038
13
38
20
28
Perimeter filtered dendrites / Nuclei count
Line
Round
1
2
3
4

**Supp. Fig S4: Additional analysis of neural differentiation in invdupdel(8p) cells.**
**(A)** Scatterplot and correlation of nuclei counts in images (ND1 & ND2) both manually counted (x-axis) and counted using CellProfiler (y-axis).
**(B)** Scatterplot and correlation of MAP2+ nuclei counts in images (ND1 & ND2) both manually counted (x-axis) and counted using CellProfiler (y-axis).
**(C)** Scatterplot and correlation of S100B+ nuclei counts in images (ND1 & ND2) both manually counted (x-axis) and counted using CellProfiler (y-axis).
**(D)** Stacked bar plot of nuclei summed across all ND rounds by label (left), and stacked bar plot of fraction of nuclei with each label (right). Colors denote label (Astrocyte, Neuron, double label, and unlabeled), x-axis separates proband and revertant line, facets indicate absence/presence of CEPT at terminal replating.
**(E)** The percentage of non-neuronal cells per image, faceted by label (astrocyte, double-label, unlabeled). Colors denote proband and revertant lines, symbols distinguish four independent rounds of differentiation. Differences in the median fraction of astrocytes, double labeled, and unlabeled cells assessed by two-tailed Wilcoxon rank-sum test (P-value above revertant boxplots relative to the proband boxplot). Medians with a significant difference in each facet are indicated below boxplots in their corresponding line color.
**(F)** Perimeter of filtered dendrites (total length of the perimeter of neurons filtered as described in the methods section divided by the number of nuclei per image. Differences in the median perimeter length of filtered dendrites / nuclei count assessed by two-tailed Wilcoxon rank-sum test (P-value above revertant boxplots relative to the proband boxplot). Colors, symbols, and medians as indicated as in Fig S4E.
